# Transient formation of collaterals contributes to the restoration of the arterial tree during cardiac regeneration in neonatal mice

**DOI:** 10.1101/2023.12.19.572474

**Authors:** Rachel Sturny, Lucie Boulgakoff, Robert G Kelly, Lucile Miquerol

## Abstract

Revascularization of ischemic myocardium following cardiac damage is an important step in cardiac regeneration. However, the mechanism of arteriogenesis has not been well described during cardiac regeneration. Here we investigated coronary artery remodeling and collateral growth during cardiac regeneration. Neonatal MI was induced by ligature of the left descending artery (LAD) in postnatal day (P) 1 or P7 pups from the *Cx40-GFP* mouse line and the arterial tree was reconstructed in 3D from images of cleared hearts collected at 1, 2, 4, 7 and 14 days after infarction. We show a rapid remodeling of the left coronary arterial tree induced by neonatal MI and the formation of numerous collateral arteries, which are transient in regenerating hearts after MI at P1 and persistent in non-regenerating hearts after MI at P7. This difference is accompanied by restoration of a perfused or a non-perfused LAD following MI at P1 or P7 respectively. Interestingly, collaterals ameliorate cardiac perfusion and drive LAD repair, and lineage tracing analysis demonstrates that the restoration of the LAD occurs by remodeling of pre-existing arterial cells independently of whether they originate in large arteries or arterioles. These results demonstrate that the restoration of the LAD artery during cardiac regeneration occurs by pruning as the rapidly forming collaterals that support perfusion of the disconnected lower LAD subsequently disappear on restoration of a unique LAD. These results highlight a rapid phase of arterial remodeling that plays an important role in vascular repair during cardiac regeneration.

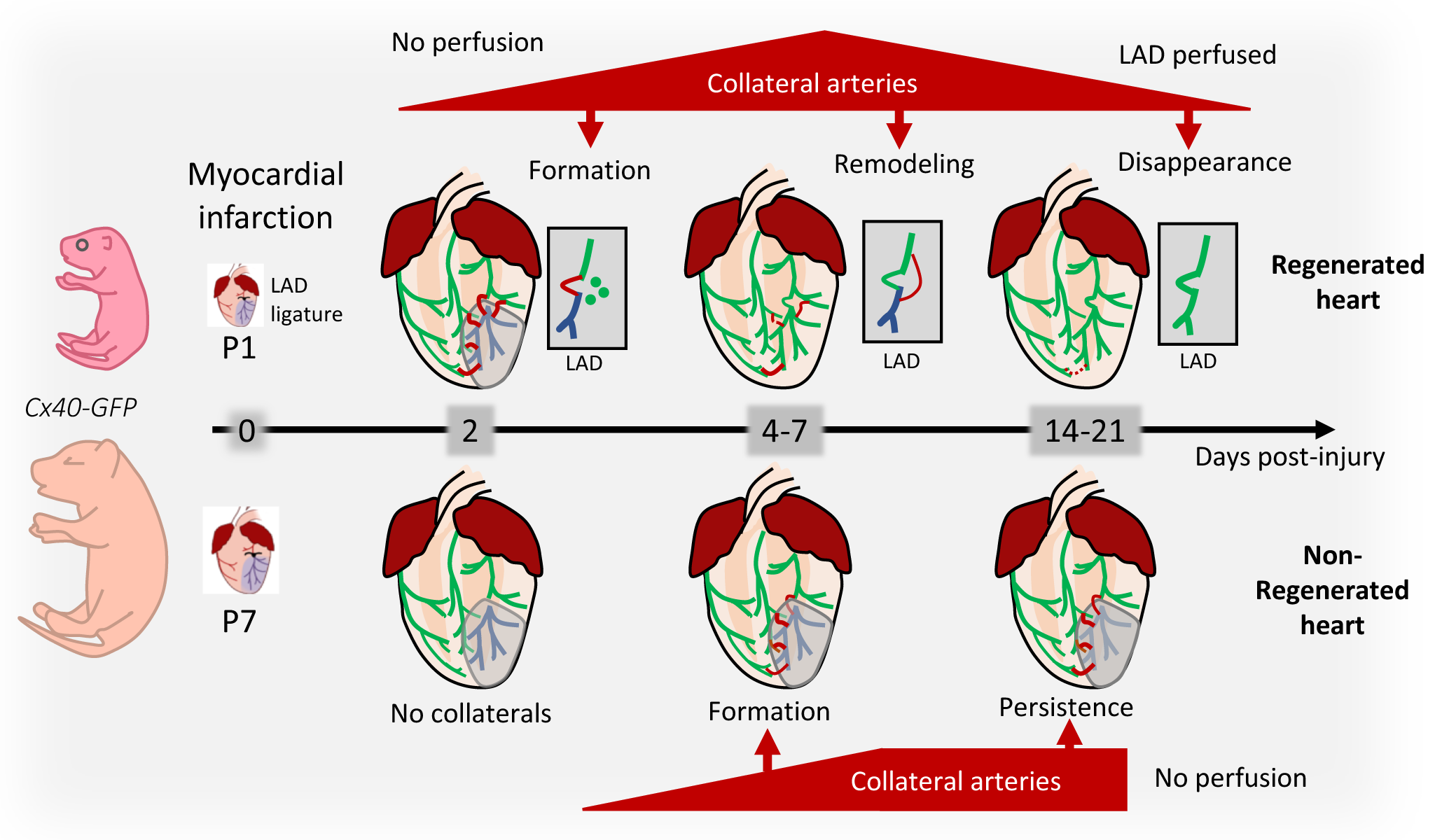

**Highlights:**

- Rapid remodeling of the arterial tree is induced by myocardial infarction.
- The endocardial response to ischemia differs between regenerated and non-regenerated hearts following myocardial infarction at postnatal days 1 or 7.
- Collateral growth is rapid and transient in regenerated hearts while delayed and persistent in non-regenerated hearts.
- Collateral arteries ameliorate cardiac perfusion and drive LAD repair.
- The restoration of the LAD occurs by remodeling of pre-existing arterial cells.

## Introduction

Cardiac ischemia due to obstruction of coronary blood flow induces cardiomyocyte (CM) death, which leads to myocardial infarction and irreversible replacement of the myocardium by fibrous scarring. The only way to save ischemic myocardium from myocardial infarction is through timely reperfusion. Myocardial infarction (MI) remains the leading cause of death worldwide, raising considerable interest in the mechanisms of blood vessel formation, with the ultimate aim of developing new and effective strategies for revascularizing the ischemic heart. The failure of clinical trials in therapeutic neovascularization through promoting angiogenesis without accompanying arterial differentiation has revealed the urgent need for a better understanding of the biology underlying cardiac revascularization [1, 2]. The blood is distributed to the myocardium through a hierarchical path composed of arteries, arterioles and capillaries. Instead of forming numerous vessels, a strategy would be to restore the arterial tree with a functional blood flow. However, such an approach requires a thorough knowledge of the remodeling of coronary arteries during myocardial infarction and a better understanding of how they develop. In the embryo, coronary arteries initially differentiate from endothelial cells within the vascular plexus giving rise to pre-artery cells [3]. Pre-specified arterial cells will migrate upstream of blood flow to build an intricate arterial network and arteriogenesis is followed by the recruitment of pericytes and smooth muscle cells [4]. In adult myocardial infarction, arteriogenesis occurs through active remodeling of pre-existing arterial cells and not from arterialization of remodeled capillaries [5]. Strikingly, we have previously shown that a major remodeling of coronary arteries occurs within the infarcted zone in the adult mouse heart associated with the formation of endocardial flowers [6]. Arterial endothelial flowers within the endocardium reveal extensive endothelial cell plasticity in the infarct zone and identify the endocardium as a site of endogenous arteriogenesis and source of endothelial cells to promote vascularization in regenerative strategies. In accordance with these results, significant endocardial remodeling was observed after myocardial infarction, reminiscent of the developmental process of coronary vessel formation taking place during trabecular compaction [7]. While the participation of endocardial cells to coronary arteries during adult myocardial infarction [8] is presumably low, the role of the endocardium following myocardial infarction in neonates is still unknown.

Under ischemia, collateral vessels represent a natural bypass to restore blood flow in the obstructed artery by forming a bridge with a healthy arteriole. The presence of well-developed collaterals has shown to be beneficial in patients with coronary artery diseases (CAD) by reducing the effects of MI and improving survival [9]. However only a third of CAD patients possess sufficient collaterals to preserve up to 25% of the normal flow and prevent MI altogether [10]. Indeed, collaterals are found in normal human hearts and in many other species with the exception of the mouse [11]. While mice develop natural collaterals in many tissues such as brain and skeletal muscle, in the heart, collaterals only appear *de novo* under ischemia [11, 12]. Recently, it has been shown that collaterals are more numerous and efficient during cardiac regeneration [13, 14]. Even if their maintenance looks advantageous, little is known about the time course of these vessels during cardiac regeneration.

Cardiac regeneration is a powerful model to identify solutions to repair the ischemic heart. The adult mammalian heart is a poorly regenerative organ because of the incapacity of cardiomyocytes to proliferate in response to damage [15]. The ability of the heart to regenerate diverges between species and, in mammals, is restricted during a short period after birth. In mice, ligation of the left anterior descending coronary artery (LAD) at birth induces a severe MI which is resolved three weeks after the surgery with restoration of normal cardiac function [15]. Strikingly, clinical reports of children who experiencing MI at birth consistently show that the patient recovered and thereafter maintained normal cardiac function suggesting that cardiac regeneration occurs also in man [16]. In zebrafish, neovascularization by angiogenesis has been recognized to precede, and possibly drive, cardiac regeneration [17]. However, these mechanisms may differ in mammals as angiogenesis is poorly developed in the mouse heart [18]. A better characterization of the process of revascularization during cardiac regeneration is required to decipher the role of endocardial cells and collateral arteries in this process.

Here, using recently developed tissue clearing technology to obtain 3D images of an entire organ [19], we analyzed the remodeling of the arterial tree during cardiac regeneration using mouse genetic and fluorescent light-sheet microscopy. We focused our study on the formation of endocardial flowers, the time course of collateral artery development and the functional restoration of the ligated left descending coronary artery (LAD). In contrast to the adult mouse heart, endocardial flowers do not appear during cardiac regeneration after neonatal MI. Neonatal mice present a rapid remodeling of coronary arteries following injury and a rapid formation of collaterals while these mechanisms are present but delayed in non-regenerating hearts. A major difference between regenerated and non-regenerated hearts, is that collaterals are respectively transient or persistent over the time course analysed. The disappearance of collaterals after regeneration is linked with the recovery of a perfused LAD after 3 weeks in all regenerated hearts. Genetic tracing demonstrate that the restoration of the LAD arises from pre-existing arterial endothelial cells independently of whether they originate in large arteries or arterioles.

## Material and Methods

### Ethics statement

All studies and procedures involving animals were in strict accordance with the recommendations of the European Community Directive (2010/63/UE) for the protection of vertebrate animals used for experimental and other scientific purposes. The project was specifically approved by the ethics committee of the IBDM SBEA and by the French Ministry of Research (APAFIS #16221-2018072015086233 v2). Husbandry, supply of animals, as well as maintenance and care of the animals in the Animal Facility of CNRS-IBDM (facility license #G-13 055 21) before and during experiments fully satisfied the animals’ needs and welfare.

### Mouse lines

The *Cx40(Gja5)-GFP* [20], *Cx40-CreERT2* [21], *R26R-YFP* [22] and *R26R-TdTomato* [23] mouse lines have been previously reported. Mice were housed at room temperature of 22 ± 3 °C, at a relative humidity between 45% and 65% and light:dark cycle of 12/12 h.

### Tam injection

For genetic tracing analysis at E14.5, *Cx40-CreERT2* males were crossed with *R26R-YFP* females and tamoxifen (150 µl) was injected intraperitoneally to pregnant females. Tamoxifen (T5648, Sigma) and progesterone (P-3972-5g, Sigma) were dissolved at the concentration of 20 mg ml−1 and 10mg.mL-1, respectively in sunflower oil. Newborn mice were recovered after caesarian and given for adoption to CD1 females. For genetic tracing analysis at E18.5, *Cx40-CreERT2* males were crossed with *R26R-tdTomato/Cx40-GFP* females and tamoxifen (150 µl) was injected intraperitoneally to pregnant females. Tamoxifen (T5648, Sigma) were dissolved at the concentration of 20 mg ml−1 in sunflower oil.

### Surgery

Myocardial infarction was induced surgically at P1 or P7 on mice by permanent ligation of the left anterior descending artery as previously described *^11^*. Sham-operated animals underwent the same surgical procedure with the exception of coronary ligation. Following analgesia with buprenorphine (0.05mg/kg), neonates were briefly sedated with isoflurane induction (chamber 5% isoflurane) followed by hypothermia-induced anesthesia on ice (4min). Neonates were then placed on an ice pad for surgery. Skin was incised, pectorals major et minor muscles were sectioned and the thorax opened in the 4th intercostal space. The left anterior descending artery was ligated just below the atria with non-resorbable 10-0 Ethilon (FG2820, Ethicon). Thorax and skin were closed with a resorbable 8-0 vicryl (V548G, Ethicon). Finally, neonates were reanimated at 37°C.

### Dissection and WGA perfusion

To avoid blood clots, mice received an intraperitoneal injection of heparin before cervical dislocation (25 USP heparin solution/g ; SIGMA-*H3393-100KU*). Hearts were then harvested, cannulated via the aorta with a blunt syringe needle, and retro-perfused with 2 to 5 mL of 1× PBS, depending of the stage of dissection. The blood was visibly flushed out from the coronary arteries under a binocular microscope. Then WGA lectin perfusion was performed with 100-200µl of 20µg/ml of Wheat Germ Agglutinin, CF®555 Conjugate (Clinisciences *29076-1) or* CF®633 Conjugate (clinisciences *29024-1*). The WGA solution was injected through the aorta, with constant manual pressure. The volume of WGA perfusion depends of the heart size. Global external views were directly acquired with a Zeiss Axio Zoom.V16 to validate the correct position of the ligation on the LAD and the WGA perfusion. Hearts were then fixed in 4% paraformaldehyde (15714 Electron Microscopy Sciences) at 4°C for 48h, and washed 3 times for 15 minutes each with 1X PBS at 4 °C under rocking .

### Tissue clearing and imaging

Tissue clearing of hearts was performed in accordance with the CUBIC protocol reported by [24] . The CUBIC-1 reagent was prepared as a mixture of 25 wt% urea, 25 wt% N,N,N0,N0-tetrakis(2-hydroxypropyl) ethylenediamine, and 15 wt% Triton X-100 in deionized water. Each fixed heart was immersed in 5ml of CUBIC-1 reagent at 37°C with gentle shaking for 1 day, after which the solution was exchanged and the sample immersed if needed in the same volume of fresh CUBIC-1 for an additional day, until imaging. All incubation steps were carried out in the dark. After imaging, the sample was washed with1X PBS, immersed in 30% (w/v) sucrose in PBS, and stored in O.C.T. compound at –80°C.

### Imaris processing

Images of cleared hearts were acquired using Lightsheet microscopy Z.1 (Carl Zeiss, Jena, Germany) equipped with a 5× objective lens (EC Plan-Neofluar 5×, numerical aperture (NA) = 0.16, working distance (WD) = 18.5 mm). Heart samples were immersed in the CUBIC-1 reagent during image acquisition. Image processing and maximum intensity projections were performed using Zen software (Carl Zeiss). Three-dimensional (3D) reconstruction of Z-stacks were conducted using Imaris software (Version 9.7, Bitplane). The entire 3D heart images was obtained from the merging of two images using IMARIS Stitcher 10.1. The IMARIS Neurofilament tracer tool was used to easily trace the arteries (Cx40-GFP^+^), with a semi-automatic autopath method. Thus, the 3D arterial tree was reconstructed, and each main artery could be color painted. The Filament selection is used to define junctions between arteries and localize collaterals.

### Antibodies

Antibodies used in this study are specific to GFP made in chick (1:500; 1020, Aves), VEGFR2 in goat (1:25; AF644, R&D system), Endoglin in rat (1:25; MJ718, DHSB), SMA-FITC in mouse (1:500; F3777, Sigma), SM22 in goat (1:100; ab10135, Abcam).

Secondary antibodies: Donkey anti-goat-647 (1:250; A21447, Life technologies), Donkey anti-chick-488 (1:250; 703 545 155, Interchim), Donkey anti-chick-647 (1:250; 703 605 155, Jackson ImmunoResearch), and Donkey anti-Rat 647nm (1:250; 712 605 153, Jackson ImmunoResearch) WGA-555 (1:500; Wheat germ agglutinin, 29076-1, Clinisciences), WGA-647 (1:500; Wheat germ agglutinin, 29024-1, Clinisciences).

### Histology

To obtain full diastole, mice received an intraperitoneal injection of 10µL/g of heparin before cervical dislocation. Hearts were then dissected out and PBS-KCl (50mM) was immediately retro perfused by the aorta. Global external views were directly acquired with a Zeiss Axio Zoom.V16 to validate the correct position of the ligation on the artery. Hearts were then fixed in 4% paraformaldehyde (15714 Electron Microscopy Sciences) overnight at 4°C, washed in PBS. Cryoprotection was achieved by successive baths of sucrose 15%, 30% and OCT (VWR Chemicals ref. 361603E) from 12 to 24h each, followed by freezing in OCT on dry ice and storage at -80°C before sectioning.

For histological studies, hearts were sectioned at a thickness of 12µm for newborn stages (P3-P5) and 20µm for later stages (P22). Sections were preserved at -80°C before immunostaining.

### Immunostaining

Sections were rinsed in PBS 1x and then permeabilized for 20 min (PBS 1×/0.2% Triton X100) and blocked for 1 h (PBS 1×/2% bovine serum albumin (w:v)/0.1% Triton X100/0.05% Saponin). The primary antibodies and WGA were incubated in blocking buffer for overnight at 4 °C. Sections were rinsed (PBS 1X/ 0.05% Tween). Secondary antibodies coupled to fluorescent molecules were incubated in blocking buffer for 1 h. After washes (PBS 1X/ 0.05% Tween), sections were mounted with DAPI-fluoromount (Ref 010-20 Southern Biotech).

### Quantifications and statistics

#### Collateral Number

The number of collateral arteries were quantified from images of whole hearts acquired using a 5X objective from whole hearts.

#### LAD statistics

To measure the development on the LAD over time in sham or MI conditions, IMARIS Statistics tools were exploited, with the filters Selected Filament and Average. Specific data were extracted : the Filament Dendrite Length (SUM), the Filament N° Dendrite Branching Points, and the Filament N° Dendrite Terminal Points. To compare the density of LAD branching vessels, Filament length was normalized for each heart height.

Statistical analyses and graphical representations were performed using GraphPad Prism 8.0.1 (Boston, Massachusetts USA). All experiments were carried out with at least three biological replicates. All statistical tests were performed using unpaired two-tailed Student’s t-test between groups with a confidence level of 95%, unless specified. Differences were considered statistically significant at p value<0.05. Data and error bars are presented as mean ± standard deviation (SD). p-values are specified in the corresponding figure when significant or close to significant.

## Results

### Early remodeling of the coronary arterial tree induced by neonatal cardiac injury

Using the *Cx40-GFP* mouse line, in which GFP is expressed in coronary artery endothelial cells, and 3D imaging of cleared hearts, we visualized the whole arterial tree and described its remodeling during cardiac regeneration (Figure 1a, b). Myocardial infarction (MI) was induced by permanent ligation of the left anterior descending artery (LAD) in mouse neonates at P1 as a model of cardiac regeneration. Coronary arterial tree remodeling was analyzed at 2, 4, 7- and 14-days post injury (dpi). Each artery was annotated using filament tracer software from IMARIS, the width of each segment indicating artery diameter (Figure 1b). Branching of the three main coronary arteries (Left, Right, Septal) was drawn using filament touch, and painted using the following color code: left=blue; right=green; septal=orange (Figure 1c). After ligation, the LAD is disrupted and divided in two parts: the upper part (light blue) and lower part (dark blue) (Figure 1c). At 14dpi, a unique LAD is rebuilt during cardiac regeneration. Our time course analysis shows an increased arterial density between 2 and 14 days in both sham and MI hearts. The density of the arterial tree was quantified for the LAD using IMARIS statistical tools in sham during the normal growth of the heart. The filament full length serves as a readout of the total length of the whole arterial tree normalized by heart height, the number of branches and end points referred to the complexity of the arterial tree (Figure 1d). These three parameters increased linearly over time in association with the physiological growth of the heart size during the same period (Figure 1d). In contrast, quantification of these parameters in infarcted hearts revealed an increased density and complexity of the LAD tree in MI compared to sham-operated hearts at 2dpi (Figure 1e, Supplementary figure 1). However, the density of the arterial tree is similar between sham and MI hearts at 4 and 7 dpi while it decreases at 14 dpi (Figure 1e). These data attest to the rapid remodeling of the LAD tree in response to neonatal MI while this phenomenon does not persist during cardiac regeneration. The slight decrease of LAD density may be explained by the remodeling of the Right and Septal arteries that invade the myocardial territory normally irrigated by the LAD.

**Figure 1:**
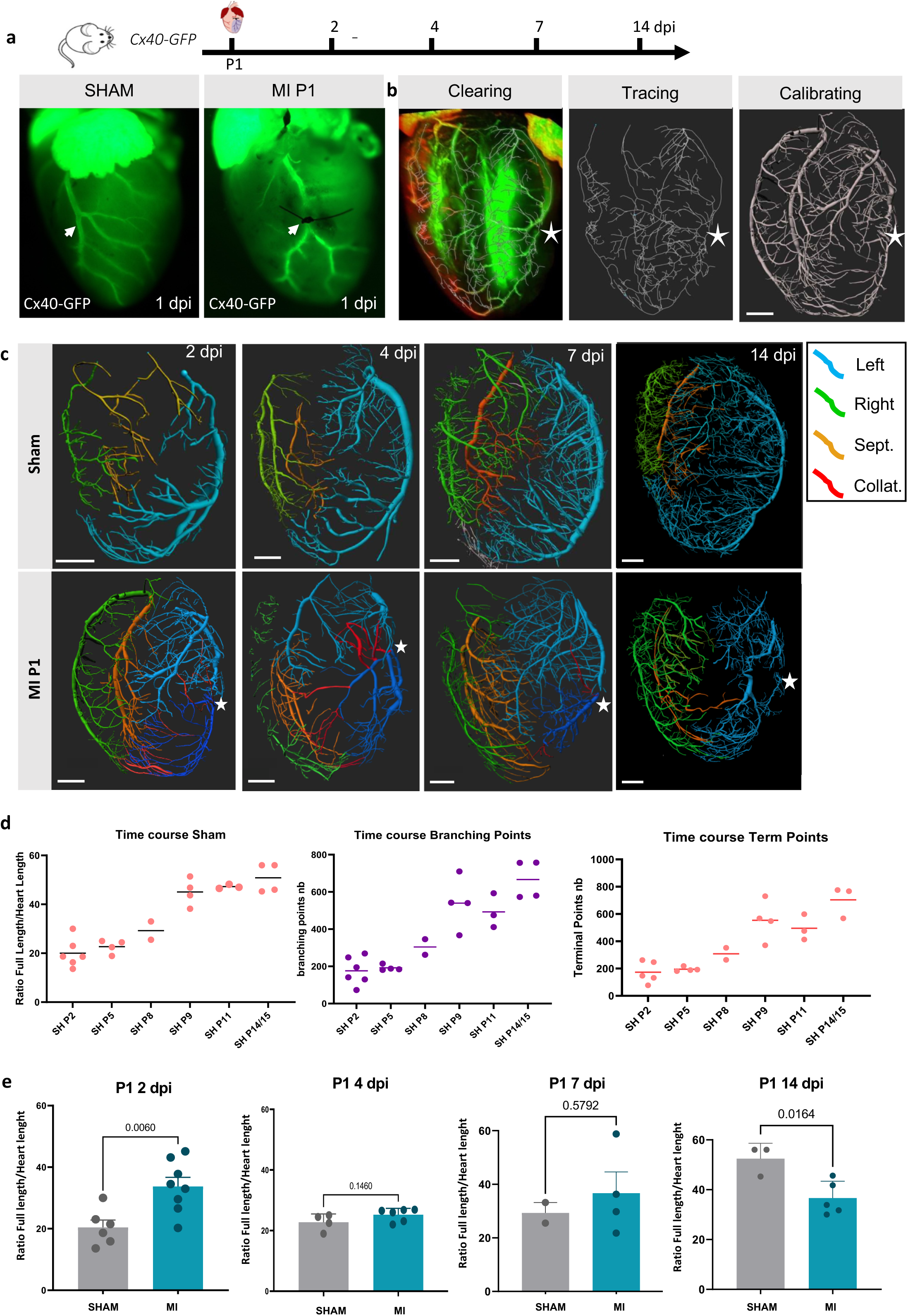
Arterial remodeling during cardiac regeneration following neonatal MI. **a**- Left lateral views of hearts from sham-operated or neonatal MI 1 day after the surgery. The position of the ligation is visible on the Connexin40-GFP-expressing coronary artery and indicate by an arrow. **b**- Example of the three procedures necessary to build the 3D image of coronary arterial tree using Imaris: clearing and imaging, filament tracing, calibration. The star indicates the ligation point. Scale bar= 500µm **c-** 3D reconstruction of the arterial tree of Sham- or MI-operated hearts at P1 and recovered at different timepoints during cardiac regeneration (2, 4, 7 or 14 days post injury). The Left coronary artery is painted in blue, the right in green, the septal in orange, and collaterals in red. The ligation point is indicated by a star. Scale bar= 500µm **d**- Time course of the density and complexity of the arterial tree in sham-operated (SH) hearts recovered from 2, 5, 8, 9, 14 and 15 postnatal day-old mice (P) (n=20). Measurements of total filament length, branch points and end points extracted from the statistics of the left coronary artery filament tracing tool of Imaris. **e**- Comparison of the arterial tree density between sham- and MI-operated hearts at 2, 4, 7 and 14 days post-injury. (t-Test, P value indicated above each graph, each dot corresponds to one heart)

Previously, we have discovered new arterial vessel formation following adult MI from the endocardium giving rise to endocardial flowers [6]. To understand if this arterial remodeling mechanism also occurs during cardiac regeneration, we investigated whether such structures are present in regenerating or non-regenerating hearts after MI induced at P1 or P7, respectively. Endocardial flowers characterized by Cx40+/VEGFR2+/Eng- staining are present in the infarct zone of non-regenerating hearts (MIP7, Supplementary figure 2A) while they were not observed following neonatal MI (MIP1, Supplementary figure 2B). Thus, these structures are not associated with cardiac regeneration. However, we observed some phenotypic changes of the endocardium during cardiac regeneration. Two days after injury, a small number of endocardial cells express a low amount of VEGFR2 but not Cx40 while they maintain high expression of endoglin (Supplementary figure 2). Seven days post-injury, we observed few GFP+ cells in the endocardium indicating that they have induced the expression of Cx40 while maintaining endoglin expression. Thus, the endocardium localized at the border of the infarct zone appears sensitive to the ischemic formation of new arteries from the endocardium.

### Collateral growth is transient in regenerated hearts and persists in non-regenerated hearts

Our 3D images of the arterial tree revealed the presence of collateral arteries which are defined as the fusion between Cx40-GFP+ ramifications coming from two main arteries and painted in red (Figure 1c, 2a, supplementary movie 1). The presence of collaterals was only detected in hearts following MI, no collaterals were observed in Sham-operated hearts (Figure 1c).

To better characterize the presence of collaterals during cardiac regeneration, we first classified collaterals in three categories based on the two main arteries that are interconnected: Right-Left (R-L); Septal-Left (S-L) or Left-Left (L-L). Right-Left collaterals are mainly present in the apex of the heart and allowed a connection with the lower part of the LAD (Figure 2a). In contrast, Septal-left junctions are present along the vertical axis of the heart with connections between the upper and/or lower parts of the LAD (Figure 2a). Moreover, numerous collaterals allow the junction between branches of the left coronary artery above and below the ligation point (Figure 2b). Our data showed that the main category of collaterals is composed of branching septal with left arterial ramifications below the ligation point (Figure 2b). Then, we quantified the number of each category of collaterals per heart during the process of cardiac regeneration at 2, 4, 7 and 14 dpi (Figure 2b, c). Collaterals are detected as early as 2 dpi, with a maximum of collaterals observed at 4 dpi. Interestingly, following MI at P1, the number of collaterals decreased after 7 dpi suggesting that this mechanism is transient during cardiac repair (Figure 2c, 2e).

**Figure 2:**
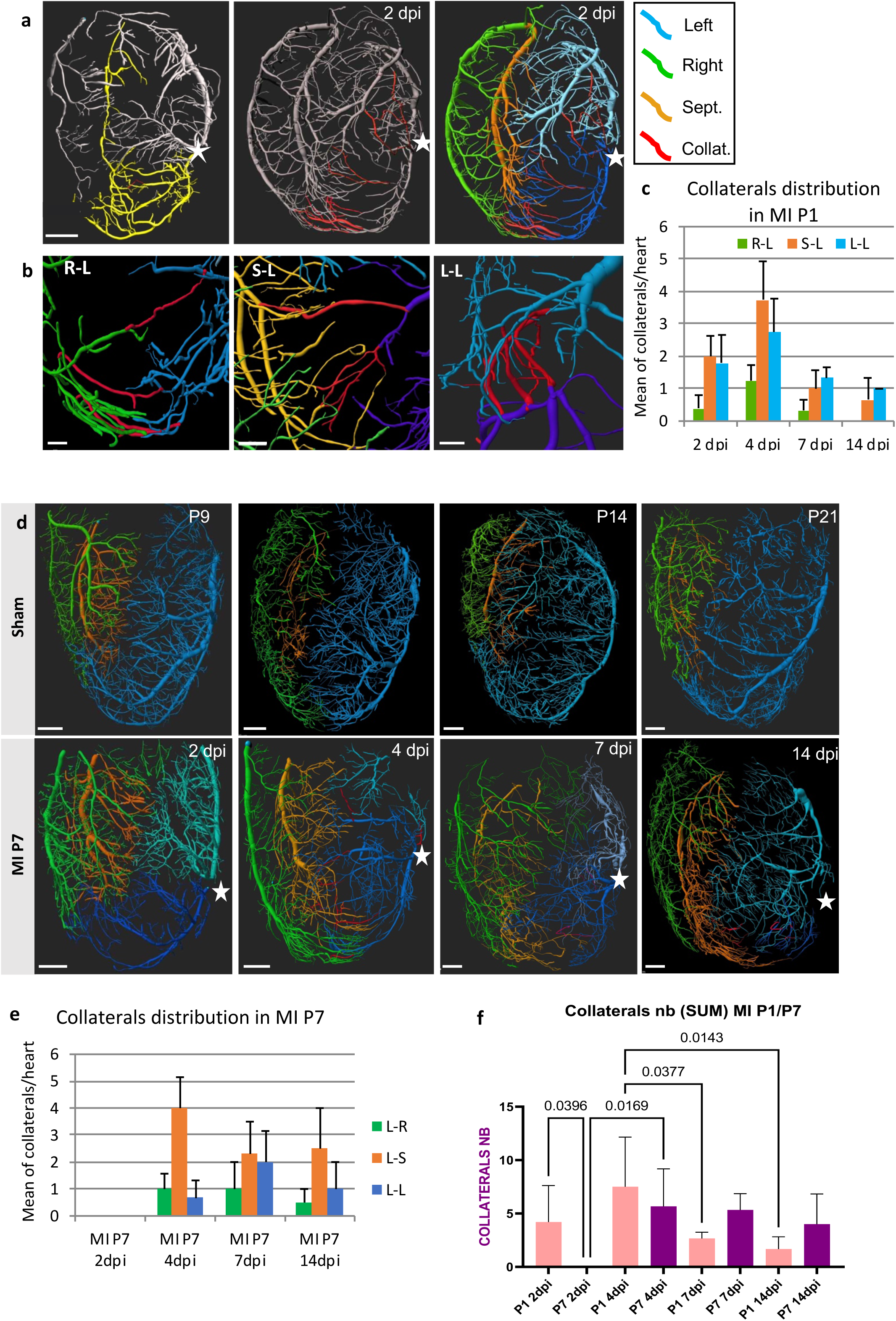
Time course of collaterals in regenerated (MI P1) and non-regenerated (MI P7) hearts. **a**- Traced collaterals on the 3D images of the arterial tree by using the Imaris filament touch tool, showing the collaterals in red and applying the color code for the other arteries (L, R and S). The star indicates the ligation point. Scale bar=500µm **b-** Examples of the three types of collaterals (Right-Left: R-L; Septal-Left: S-L; Left-Left: L-L). Scale bar=100µm **c-** Quantification of the mean of each type of collaterals per heart during the time course of cardiac regeneration (2, 4, 7 and 14 days post-injury; N=3 per stage). **d-** 3D reconstruction of the arterial tree of Sham- or MI-operated hearts at P7 and analyzed at different timepoints after injury (2, 4, 7 or 14 days post-injury). The Left coronary artery is painted in blue, the right in green, the septal in orange, and collaterals in red. The ligation point is indicated by a star. Scale bar= 500µm **e-** Quantification of the mean of each type of collaterals per heart during the time course after injury (2, 4, 7 and 14 days post-injury; N=3 per stage). **f-** Comparison of the total number of collaterals per heart in regenerated (MIP1) versus non-regenerated (MIP7) hearts during the time course after the injury (2, 4, 7, and 14 days post-injury). (One way Anova, Fisher test, P value indicated above each graph, N=5 P1 2dpi; N=4 P7 2dpi, P1 4dpi; N=3 P1 7-, 14-dpi, P7 4-,7-,14-dpi)

To understand whether the development of collaterals is specific to regenerated hearts, we performed the same experiments on 7-day-old mice, in which myocardium is non-regenerative (Figure 2d). In contrast to MI P1 hearts, the density of the left arterial tree decreases after surgery and tends to remain low in comparison to sham hearts (Supplementary figure 3). This is compensated by an extension of the right or septal coronary arteries colonizing the apex of the heart (Figure 2d). Collaterals are absent in hearts at 2dpi but present from 4dpi and persist up to 14dpi (Figure 2d, e). The distribution of the different categories of collaterals with a majority of S-L type is similar to that observed in MI at P1. Thus, the presence of collaterals in non-regenerated hearts suggests that remodeling of coronary arteries is not a mechanism exclusively limited to regenerating hearts. However, in contrast to MI P1 hearts, the presence of numerous collaterals persists at 14dpi after the scar has formed (Figure 2f). To explain this discrepancy between MIP1 and MIP7, we asked whether the restoration of the LAD as observed above could replace collaterals during regeneration and would therefore only occur in regenerated hearts.

### Complete repair of the LAD following myocardial infarction is specific to regenerated hearts

To understand whether the restoration of the LAD is specific to regenerated hearts, we compared the morphogenesis of the LAD at different timepoints: 1, 2, 4, 7 and 14 days after injury. A close look at the GFP staining at the level of the ligature demonstrates the dynamic evolution of the LAD which is disrupted at day 1 and 2 and then the upper and lower parts are reconnected by a small arterial vessel which remodeled with time to reach its normal size at 14 dpi (Figure 3a). A deep analysis of 3D images using Imaris shows the presence of numerous collaterals in this region at 4 dpi in MIP1 while a principal connection of the upper and lower parts of the LAD is detected from 7 dpi only (Figure 3b). This suggests that arterioles remodeled faster than main arteries. Interestingly, the restoration of the LAD was observed in only one MIP7 heart at 14dpi while collaterals are observed in this region in all MIP7 hearts from 4 dpi and persist at least until 14dpi (Figure 3b, 3e). However, the restoration of a large LAD is detected in <15% of MIP7 operated-hearts versus >90% of the MIP1 hearts (Figure 3d). This suggests that the restoration of the LAD appears independently of the presence of collaterals. In the only MIP7 heart whose LAD was restored, the presence of a large scar under the ligature attests to a lack of regeneration (Figure 3c). Thus, the restoration of the main LAD is not sufficient for cardiac regeneration.

**Figure 3:**
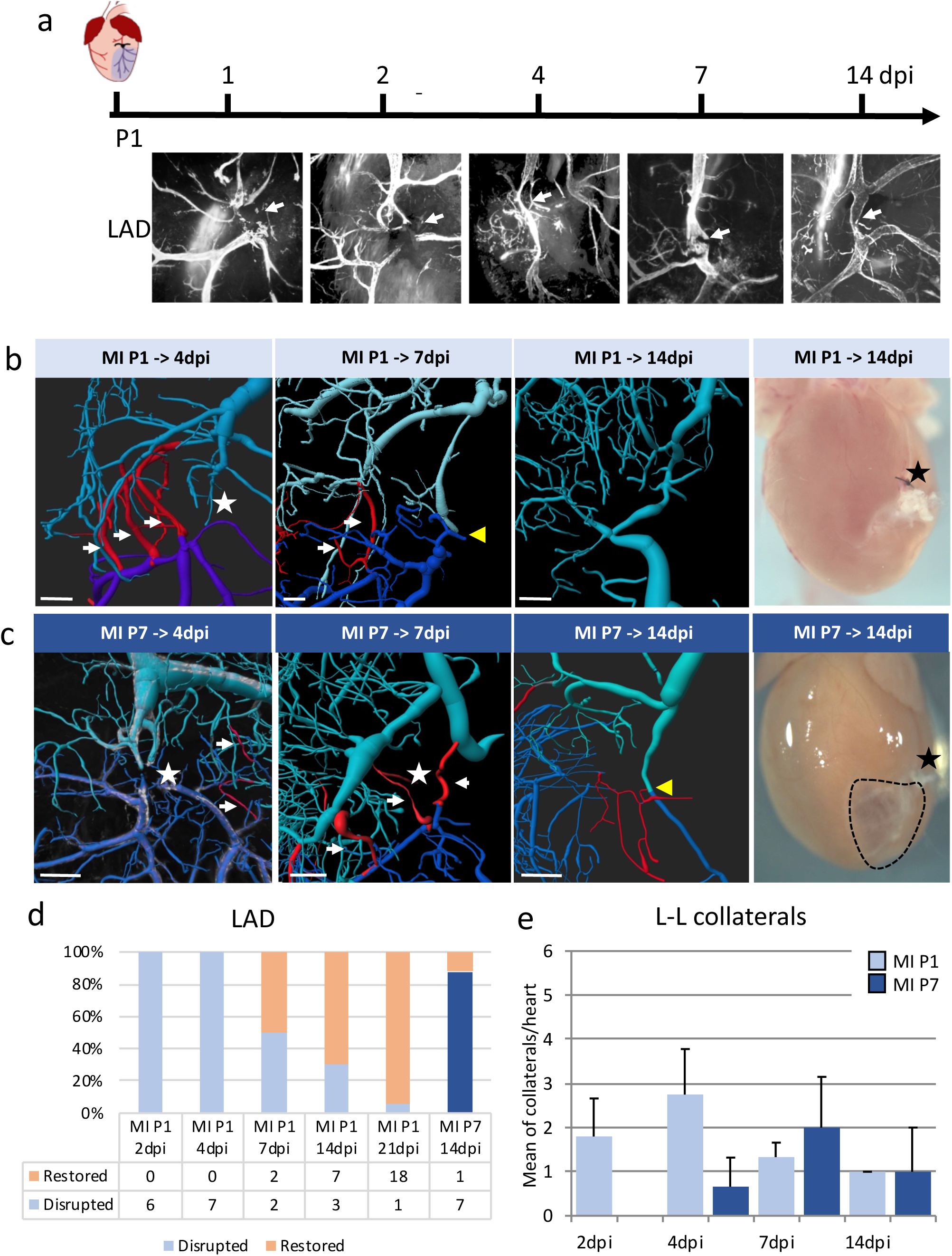
Time course of restoration of the left coronary artery in regenerated (MI P1) and non-regenerated (MI P7) hearts. **a-** Maximum intensity projection of the Left coronary artery at the ligation point (white arrow) of Cx40-GFP MIP1 hearts during the time course of cardiac regeneration (1, 2, 4, 7, and 14 days post-injury). **b-** 3D-reconstruction of the collaterals (white arrows) and the left coronary artery (yellow arrowhead) at the ligation point (star) of MI-operated hearts at P1 and recovered at different timepoints after the injury (4, 7 or 14 days post-injury). The right panel is a picture of a regenerated heart 14 days post-injury with a knot on the LAD. Scale Bar= 100µm **c-** 3D-reconstruction of the collaterals (white arrows) and the left coronary artery (yellow arrowhead) at the ligation point (star) of MI-operated hearts at P7 and recovered at different timepoints after the injury (4, 7 or 14 days post-injury). The right panel is a picture of a non-regenerated heart 14 days post-injury with a knot on the LAD and the fibrotic scar below (dotted line). Scale Bar= 100µm **d-** Quantification of the number of hearts with disrupted or restored LAD during the time course of cardiac regeneration (2, 4, 7, 14, and 21 days post-injury). **e-** Comparison of the mean number of L-L collaterals per heart in regenerated (MIP1) versus non-regenerated (MIP7) hearts during the time course after injury (2, 4, 7, and 14 days post-injury).

### Collateral artery growth ameliorates cardiac perfusion and drives LAD repair

Next, we asked when the remodeling of the arterial tree following MI at P1 is functional by studying the passive perfusion of these vessels. To do so, a solution of WGA lectin was retro-injected through the aortae to label perfused arteries. The volume injected was adjusted to label the arterial tree in sham-operated hearts similarly between 2 to 21 dpi (Figure 4a). The WGA staining comprises Cx40-GFP-positive vessels including main arteries and secondary arterioles, as well as Cx40-GFP-negative vessels, but not capillaries or veins (Figure 4a). The same volumes were used to label the arterial tree in MI hearts at 2, 4, 14 and 21 dpi. While Cx40-GFP staining persists after the injury, WGA staining was not detected in the disrupted artery below the ligation, showing inefficient perfusion of the infarcted area at 2 and 4 dpi (yellow arrowheads in Figure 4b). In the lower LAD we observed perfusion in a part of disconnected arteries in association with the presence of functional collaterals (red arrowheads in Figure 4b). Interestingly, in another heart at 2 dpi, WGA staining was detected in a vessel connecting the upper and the lower LAD while this vessel is Cx40-GFP negative suggesting that perfusion may precede arterial differentiation during the process of collateral formation (Figure 4c). At 7 dpi, collaterals and the LAD are both functional while only a large perfused LAD is detected at 14 dpi and 21 dpi (Figure 4b). At 21 dpi, the restored LAD presents two types of trajectory suggesting different mode of repair. First, we observed a tortuous vessel with a characteristic S-shape at the ligation point (type I), consistent with a repair of the LAD through collaterals localized on the trajectory of the previous LAD. The second type of restored LAD shows a branch interrupted to the right and a LAD shifted to the left (type II) in line with a repair of the LAD through collaterals localized between the lower LAD and a lateral arterial branch. These data demonstrate that restoration of the LAD during cardiac regeneration is not stereotypical but the process requires the rapid formation of collaterals to support perfusion of the disconnected lower LAD which disappeared on restoration of a unique LAD. Our data suggest that the trajectory of the restored LAD depends on the localization of perfused collaterals.

**Figure 4:**
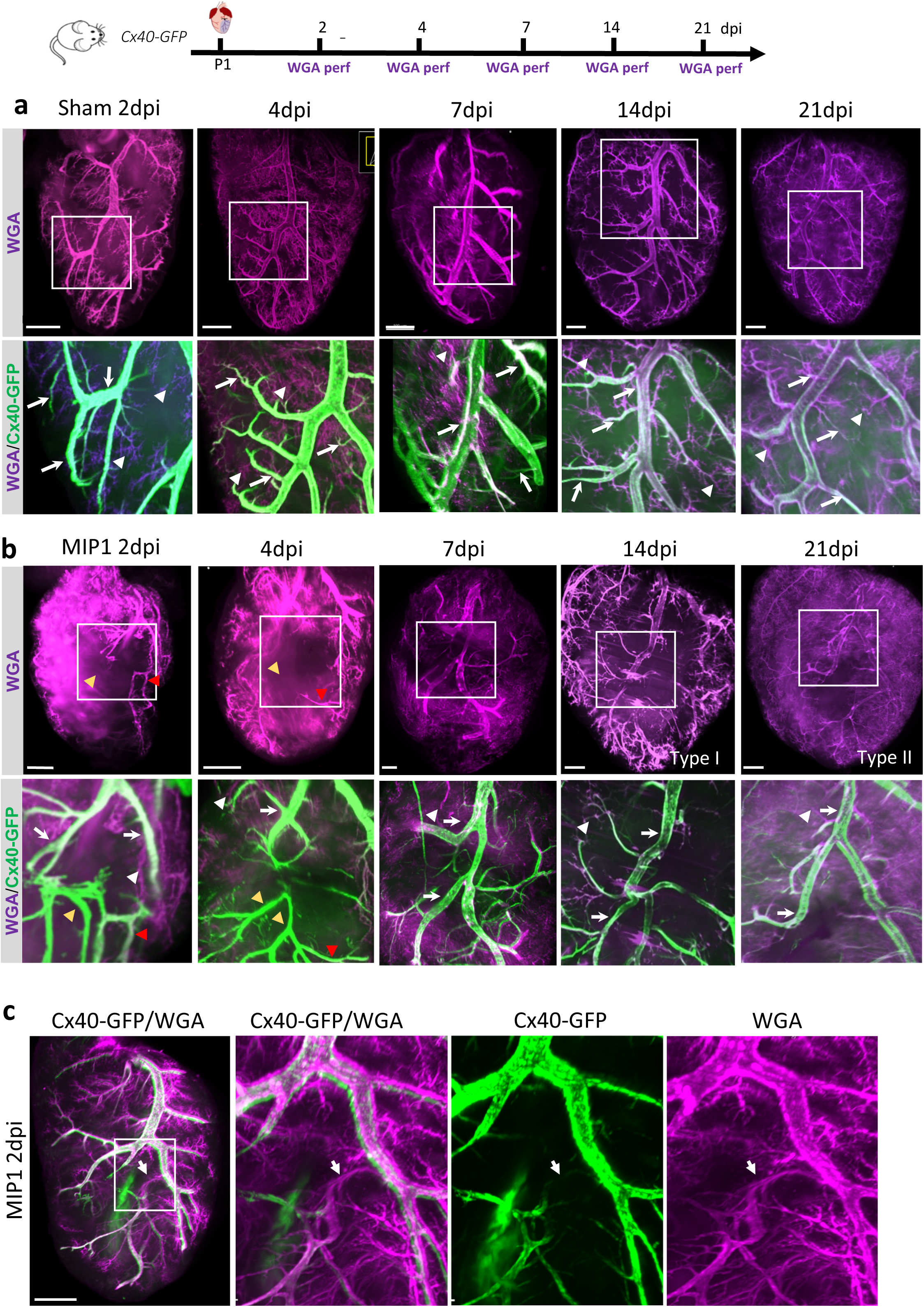
WGA lectin perfusion during the time course of cardiac regeneration. **a-** Maximum intensity projection of WGA-perfusion (Purple) of the Left coronary artery in sham-operated heart at different timepoints after the injury (2, 4, 7, 14, and 21 days post-injury). Below are high-magnification images of the LAD co-labelled with Cx40-GFP. Perfused Cx40-GFP arteries and arterioles are indicated by arrows while perfused small Cx40-negative vessels are indicated by arrowheads. Scale Bar= 500µm **b-** Maximum intensity projection of WGA-perfusion of the Left coronary artery in MI P1-operated heart at different timepoints during cardiac regeneration (2, 4, 7, 14, and 21 days post-injury). Below are high-magnification images of the LAD at the ligation point of Cx40-GFP MIP1 hearts during the time course of cardiac regeneration. Non-perfused Cx40-GFP arteries are indicated by orange arrowheads and collateral perfusion of Cx40-positive vessels indicated by red arrowheads. Scale Bar= 500µm **c-** Example of a WGA-perfused L-L collateral 2 days after injury in absence of Cx40-GFP co-labeling. Scale Bar= 500µm

### The repair of the LAD occurs by the remodeling of arterial endothelial cells

From the literature, large collaterals at the level of the ligature are thought to develop by arteriogenesis [5]. However, this assumption was concluded from a genetic tracing analysis of capillaries on sections, where it is difficult to clearly identify collaterals at the level of the repaired artery [5]. Das et al. have described a new mechanism, i.e. artery reassembly, in which endothelial cells at artery tips migrate on capillaries to develop collateral arteries in the watershed region [14]. Our data suggest that arterioles remodeled faster than main large arteries. To better understand the mechanism of LAD repair, we asked whether the restored LAD is rebuilt from arterial cells of the main arteries or arterioles. Arterioles are built after birth while main and secondary arteries are built during embryonic development [25], so tamoxifen induction of Cx40-CreERT2 before birth will label the main artery while postnatal induction will label all arterial cells. To determine the origin of the endothelial cells contributing to the LAD repair following MI at P1, we performed a genetic tracing analysis of coronary arterial cells using *Cx40-CreERT2* crossed with *R26-dTtomato* mice and assessed lineage contributions in whole-mount hearts. First, we injected tamoxifen at E14.5 to label the main coronary arteries and compared the distribution of Cx40-derived tdTomato cells with that of Cx40-GFP arterial cells. (Figure 5a). As Cx40 is expressed in cardiac trabeculae at E14.5, a large proportion of tdTom^+^ cells are ventricular cardiomyocytes, however, tdTom^+^ cells are also present in the main LAD and secondary arteries in sham-operated hearts while absent from arterioles (white arrowheads in Figure 5b). MI was then performed at P1 and 21 days after MI we observed that the restored Type I-LAD is mainly composed of tdTom^+^ cells while secondary vessels are tdTom^-^ suggesting that the endothelial cells of the main LAD are the ones contributing to the repair. These results are consistent with the observation that isolated Cx40-GFP endothelial cells are scattered around the ligation 1 and 2 days after the surgery, suggesting rapid arterial remodeling as has been described for artery reassembly (Figure 3a and Supplementary figure 5). Type II-LAD that are thought to arise from lateral collaterals, are composed in part by tdTom^-^ cells suggesting that the recruitment of endothelial cells required for the LAD repair occurs outside the main and secondary arteries. These results were confirmed on transverse sections using *R26-YFP* reporter mice and double SMA/RFP immunostaining to label arteries. The quantification of YFP^+^/total arterial endothelium (Cx40-RFP+) coimmunostaining shows a reduced proportion of YFP^+^ of the arteries at the level of the ligation (Supplementary figure 4). The presence of new arteries formed by Cx40-negative cells suggest a contribution of other cell types than those labeling the main arteries during development. To understand if these cells originate from pre-existing arteries at the time of surgery, we performed a second genetic tracing by injecting Tamoxifen at P0, one day before the surgery allowing the labeling of all arterial cells including those in arterioles (Figure 5c). In all cases, the restored LAD and secondary vessels were tdTom^+^ indicating that ischemic-induced arterial remodeling involved only arterial cells as described for arteriogenesis and artery reassembly. Our results demonstrate the high plasticity of pre-existing arterial cells independently of whether they originate in large arteries or arterioles.

**Figure 5:**
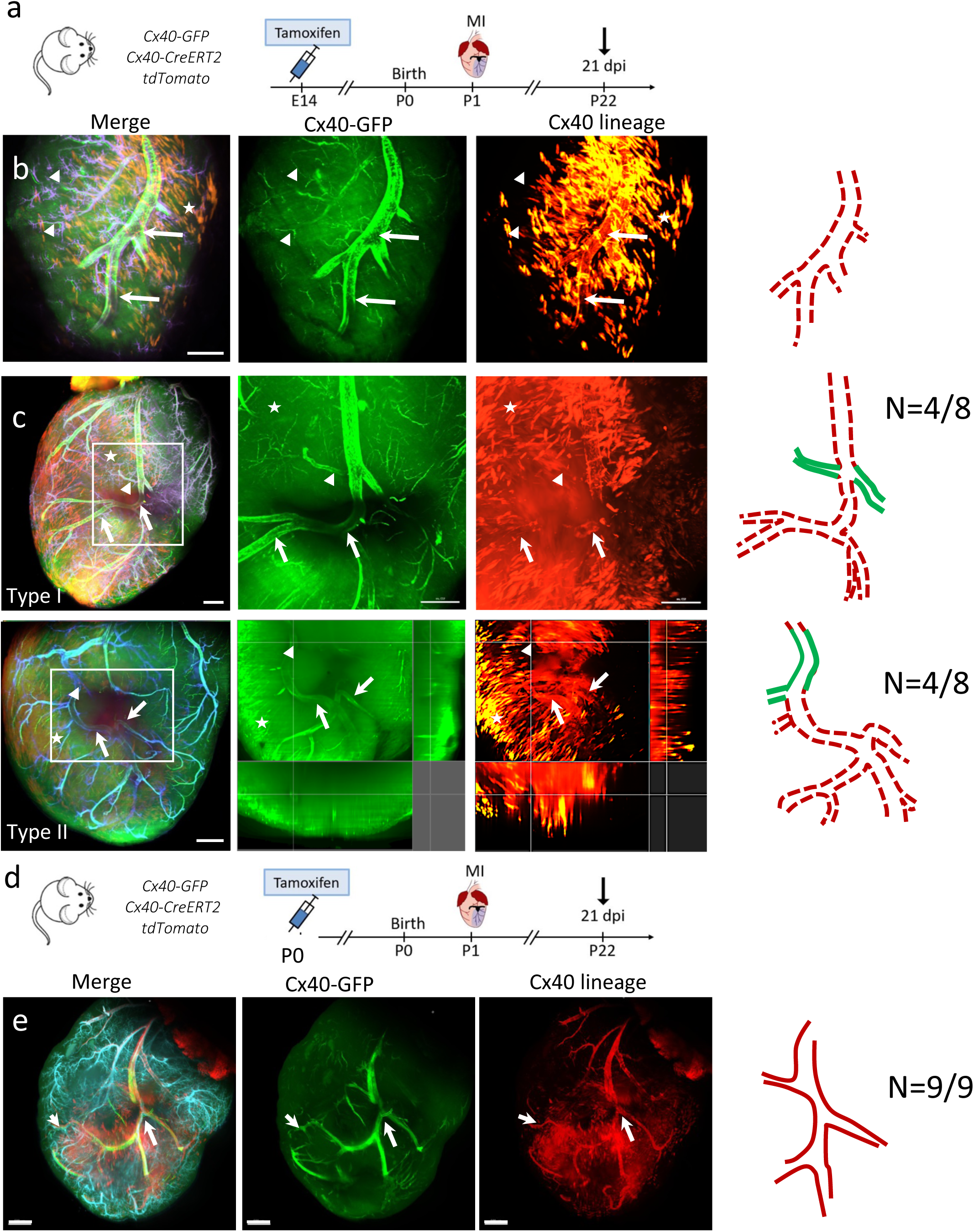
Genetic tracing of Cx40-derived arterial cells during cardiac regeneration. **a-** Experimental workflow describing the genetic tracing of Cx40-arterial cells of main arteries following tamoxifen injection at E14.5. **b-** Maximum intensity projection of a representative Left coronary artery from a sham-operated heart 21 days after the injury with the arterial tree in green (*Cx40-GFP*), Cx40-derived cells in red (*Tomato*-lineage) and WGA perfusion in blue. Main arteries are labeled in red (arrows) while arterioles are unlabeled (arrowheads). In the right panel, drawing of the LCA painted in red as derived from *Cx40-*lineage. Scale Bar= 00µm **c-** Maximum intensity projection of representative *CX40-GFP/Tomato*-lineage Left coronary artery from 2 MI-operated hearts 21 days after the injury. The main repaired LAD artery is labeled in red (arrows) in type I repair while the main LAD is unlabeled (arrowheads) in type II. In the right panel, drawing of the LCA painted in red or green as derived from *Cx40*-lineage or not, respectively. Scale Bar= 500µm **d-** Experimental workflow describing the genetic tracing of Cx40-arterial cells of main arteries and arterioles following tamoxifen injection at P0. **e-** Maximum intensity projection of representative *CX40-GFP/Tomato*-lineage Left coronary artery from a MI-operated heart 21 days after the injury. Main LAD and arterioles are indicated by arrows. In the right panel, drawing of the LCA painted in red as derived from *Cx40-*lineage. Scale Bar= 500µm

## Discussion

Revascularization of the ischemic heart is a prerequisite for cardiac regeneration. In this study, we show the important role of collateral artery formation for repair of the arterial tree and timely perfusion of the infarcted zone. However, these collaterals disappeared after the occluded LAD was completely restored and functional. Moreover, genetic tracing experiments demonstrated that the restored LAD is formed from arterial endothelial cells independently of whether they originate in large arteries or arterioles.

Knowing the importance of coronary arterial remodeling following ischemia, we developed a whole-mount imaging strategy to provide a comprehensive assessment of these events during cardiac regeneration or scar formation. We performed tissue clearing using CUBIC techniques to visualize endogenous fluorescent markers such as GFP and TdTomato in order to avoid bias from whole-mount immunofluorescent labelling which is technically challenging in post-natal hearts *^14^*. This powerful clearing technology permits the generation of 3D images of the entire arterial tree using Imaris software. We reconstructed the entire left coronary branch in multiple hearts from neonates to 3 weeks-old mice. Quantification of total vessel length, branching points and tips revealed a linear increase of the density and complexity of the arterial tree associated with the growth of the heart during this period. While this result is consistent with previous data [13, 14], our data provide a first time-course of the entire left coronary artery during this process allowing comparison with hearts following MI during regenerative and non-regenerative time-windows. Our study shows that upon ischemia at birth, the left arterial tree grows faster than in sham-operated hearts while this early response is not observed in non-regenerative hearts. However, arterial remodeling is effective a few days following the surgery in non-regenerated hearts as depicted by the invasion of the apex by the right and the septal coronary arteries, and the presence of endocardial flowers and collaterals. The difference in the timing of arterial remodeling suggests that the early response in regenerative hearts may reduce the irreversible effects of ischemia by restoring more rapid reperfusion, thus protecting the myocardium from death. In adult human hearts, it has been demonstrated that early perfusion of the obstructed artery is beneficial, however is not sufficient to avoid myocardial damages [26].

It is not only a question of timing of arterial remodeling but also of the type of response. Indeed, endocardial flowers, which are present in the infarct zone of adult [6] and P7 hearts, are absent in neonatal hearts after MI. This discrepancy may be explained by changes in the endocardium and associated cell types that do not respond to the same stimuli at P1 or P7. The appearance of endocardial flowers is preceded by a rapid expression of VEGFR2 in the endocardium following MI in response to ischemia. We observed the same response following neonatal MI, however as perfusion restarts in this region, VEGFR2 expression disappears 7 days post injury. Thus, the time course of these flowers with an appearance at 7dpi is not compatible with cardiac regeneration. Together, these data suggest that endocardial flowers do not contribute to a repair mechanism but rather accompany irreversible scar formation. In accordance with these observations, it has been shown that there is an absence of endothelial cells derived from the endocardium after adult MI [8], suggesting that the endocardial remodeling in MI does not persist and does not actively participate in the coronary vasculature post-MI. To date, these flowers have not been described in another model of repair or regeneration, however spontaneous coronary fistula resembling endocardial flowers have been seen in human hearts following myocardial infarction which are thought to arise from aberrant paths of collaterals and to spontaneously disappear [27, 28].

Our 3D images of the entire arterial tree using Cx40-GFP reporter mice highlighted the presence of deep collaterals between septal, right and left coronary branches with left obstructed coronary branches allowing effective perfusion of these territories. Similar collateral junctions have been recently identified using similar clearing techniques and whole-mount immunofluorescence and smooth muscle actin (SMA) labeling [13]. It is interesting to notice that numerous collaterals connected the septal coronary artery with the left coronary artery in similar proportion in both studies. Anbazhakan et al observed numerous, larger and more efficient collaterals in neonates than in adult hearts while we found a similar total number of collaterals in both cases [13]. However, their observations were carried out at a single time point while we performed a time-course analysis over 14 days post-injury. Our data showed that the presence of collaterals is transient in regenerated hearts whereas they persist in non-regenerated ones. This discrepancy may arise from the presence of inefficient collaterals in non-regenerated hearts [13].

Our time course shows that collaterals appeared as early as two days post-surgery in neonatal mice as has been described for adult mice [11] while we observed a delay in the formation of collaterals in non-regenerating hearts at P7. This difference can be explained by the use of different imaging techniques, namely 3D images of fluorescent arterial marker versus arterial angiography following infusion of Microfil. In our study, the detection of collaterals requires the expression of Cx40 in both vessels forming the junction, whereas using arterial angiography, the detection of collaterals depends on the formation of a lumen size sufficient to allow diffusion of Microfil. However, both studies show that a maximum number of collaterals is observed at 7dpi in non-regenerating hearts. Thus, together these data show that collateral formation is not restricted to regenerating hearts. In fact, the formation of collaterals is temporally regulated during cardiac regeneration and progressively disappears when myocardial and LAD repair occurs to completely disappear when cardiac regeneration is complete (3 weeks after surgery). In adult hearts, collaterals are still present 28 days after injury, however, these vessels are less functional than in neonates [11, 13]. Transient collateral arteries have been observed in other systems such as during reperfusion of the inferior leg where artery disappearance is known as pruning [29]. The pruning mechanism consists in the formation of a smaller number of larger vessels replacing numerous small vessels that are believed to be more performant for blood perfusion [30].

For the first time, this study looks at the restoration of the LAD during cardiac regeneration. Our time course reveals that the LAD is completely restored 3 weeks after neonatal infarction while it is restored only in 1 out of 7 non-regenerated hearts. This restoration is preceded by the development of numerous left-left collaterals at the level of the ligation forming a small junctional vessel which increase in size while the number of collaterals disappears. The formation of a large caliber vessel by pruning is correlated with the idea that collaterals present higher blood flow performance in regenerated hearts compared to non-regenerated ones [13]. Although we observed that the neo-formed LAD follows two stereotypical trajectories depending on the localization of collaterals. Type I and II may also arise from differences in the arterial tree morphology before MI, as the coronary arterial tree presents diverse patterns in the mouse [31].

Our data suggest that perfusion of collaterals occurs before the expression of the arterial marker Cx40. During development, coronary arteries form through remodeling of an immature vascular plexus in a process triggered and shaped by blood flow [32]. Indeed, Cx40 expression is directly induced by blood flow [33]. These data suggest that collaterals arteries are formed by Cx40-negative cells that express this marker once the blood flow is established. In the heart, Cx40 is specifically expressed in arterial cells and not observed in capillaries, veins and lymphatics vessels, suggesting that endothelial cells forming coronaries may arise from these types of endothelial cells. However, Das and collaborators have demonstrated, very recently, that collateral growth starts by a process of arterial dedifferentiation which could explain the absence of Cx40 expression in the early collaterals [34]. Indeed, MI-induced proliferating arterial cells (EdU+) express a low level of Cx40 compared to arterial cells in a well-formed vessel [34].

In the literature, it is unclear whether large collaterals at the level of the ligation point and collaterals of the watershed region develop by distinct mechanisms, i.e. arteriogenesis vs artery reassembly [5, 14]. However, both mechanisms implicate the formation of new arteries from arterial endothelial cells. Our genetic tracing analyses using the *Cx40-CreERT2* mouse line demonstrated that cells repairing the LAD originate only from arterial cells coming from large arteries and arterioles born, respectively, at early or late timepoints during development. These observations suggest that the repair mechanism is not dependent on the origin of the arterial cells but rather on the location of the collaterals. The process of artery reassembly proposed by Das and colleagues[14] includes the disassembly of artery tips, followed by the migration of these cells that then proliferate and coalesce de novo to form a new vessel. The presence of individual endothelial cells around the ligated LAD one day after the surgery suggest that the mechanism of artery reassembly may also be used to restore the LAD and is not restricted to distal arterioles of the watershed region. In conclusion our results provide new insights into the mechanisms by which the coronary arterial tree is restored during cardiac regeneration.

## Acknowledgments

We are grateful to Dr Gaetano D’Amato for his careful reading of the manuscript and his suggestions. We thank Brice Detailleur and Elsa Castellani for technical assistance in tissue preparation and light-sheet imaging and the Turing Centre for Living systems for image analysis. The France-BioImaging infrastructure is supported by the ANR (ANR-10-INBS-04-01, “Investissements d’Avenir”). This work was supported by the Centre National de la Recherche Scientifique (CNRS), by a grant from the Association Française contre les Myopathies (AFM). LB is recipient of a doctoral fellowship from the Ecole Normale Superieure (ENS) and of a doctoral fellowship extension from the Institut Marseille Maladies Rares (MarMaRa).

AI-assisted technology was not used in the preparation of this work.

## Author CRediT statement

**RS**: Conceptualization, Data curation, Methodology, Investigation, Validation, Visualization and Writing – Review & Editing. **LB**: Conceptualization, Data curation, Methodology, Investigation, Validation and Visualization and Writing – Review & Editing. **RK**: writing – Review & Editing. **LM**: Conceptualization, Methodology, Validation, Visualization, Funding Acquisition, Project Administration, Supervision, Writing – Original Draft Preparation and writing – Review & Editing.

---

**Supplementary figure 1:**
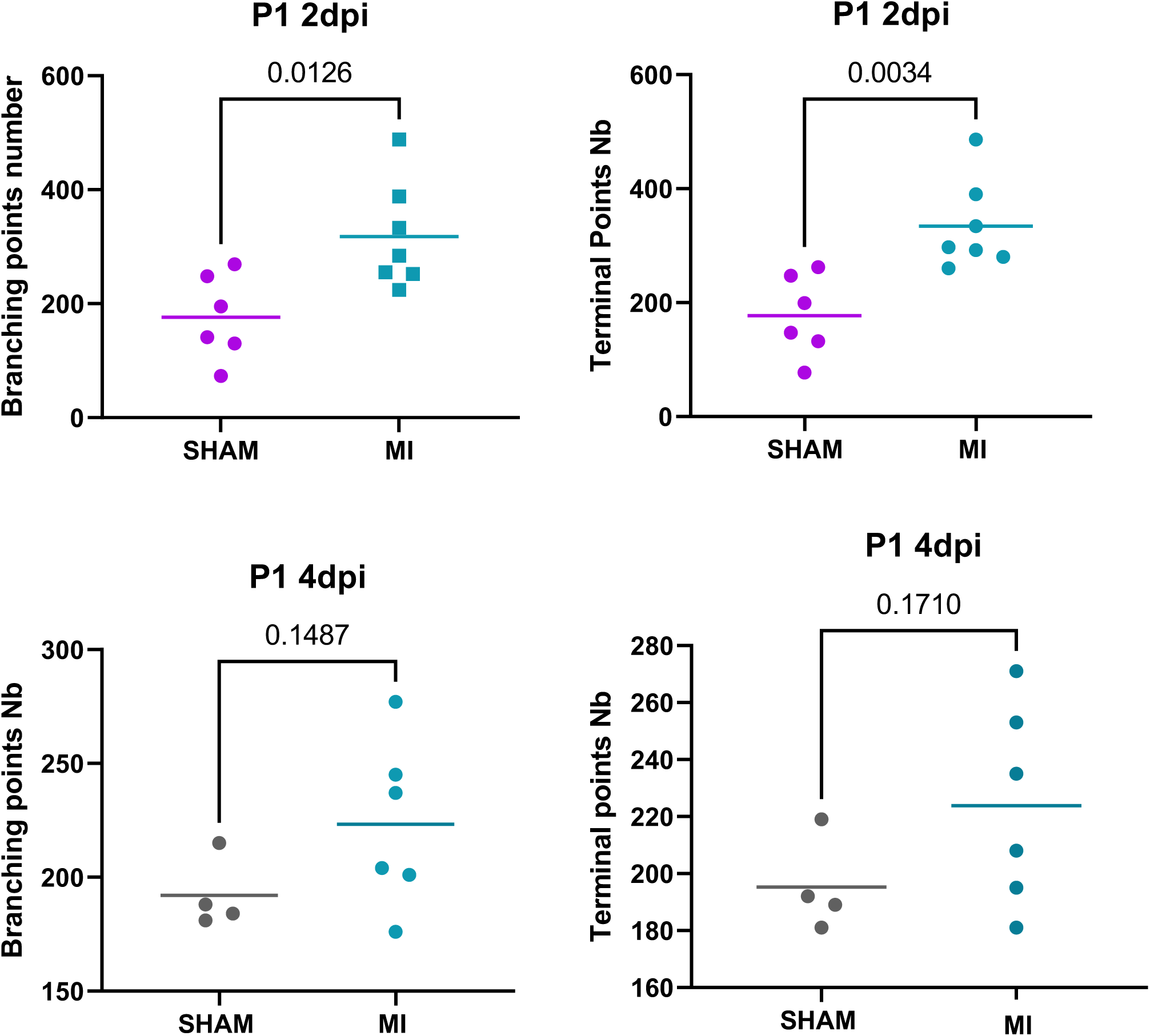
Comparison of the arterial tree complexity (branch points and end points) between sham- and MIP1-operated hearts at 2 and 4 days post-injury. (t test, p value above each graph, N: each dot corresponds to one heart)

**Supplementary figure 2:**
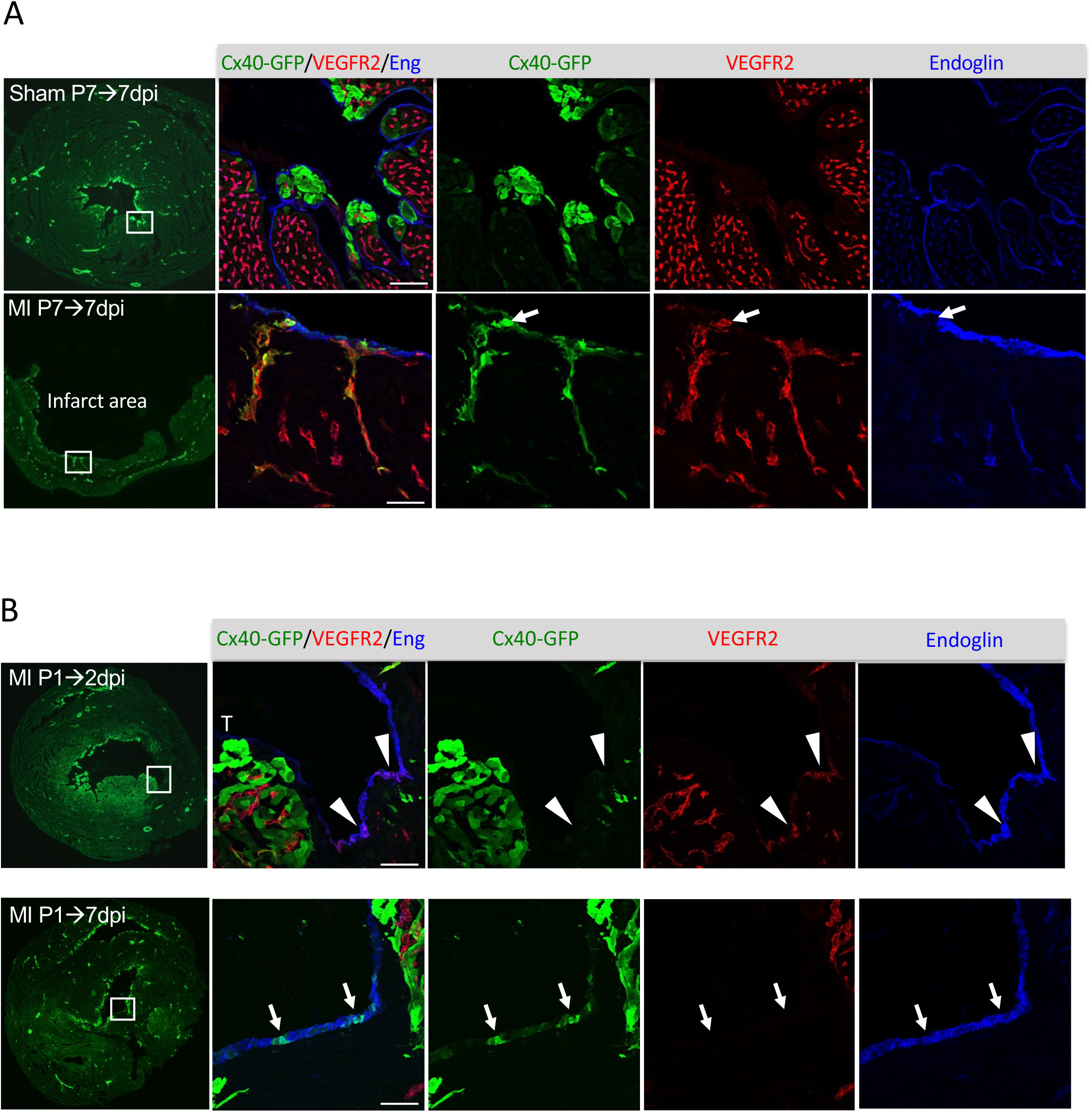
Endocardial remodeling in regenerative or non-regenerative hearts. **a-** Transverse sections of left ventricle from *Cx40-GFP* Sham and MIP7 7dpi. High magnification of the subendocardial region stained by Endoglin (endocardium), Cx40-GFP (arteries) and VEGFR2 (capillaries and endothelial progenitors). White arrows indicate the presence of endocardial flowers at the endocardial surface of the infarct area in MIP7 hearts. **b-** Transverse sections of left ventricle from Cx40-GFP MIP1 2 and 7 days post-injury. High magnification of the subendocardial region stained by Endoglin (endocardium), Cx40-GFP (arteries) and VEGFR2 (capillaries and endothelial progenitors). White arrowheads indicate the presence of VEGFR2-endocardial cells of the infarct area in MIP1 hearts 2 days post-injury. Scale bar= 50µm.

**Supplementary figure 3:**
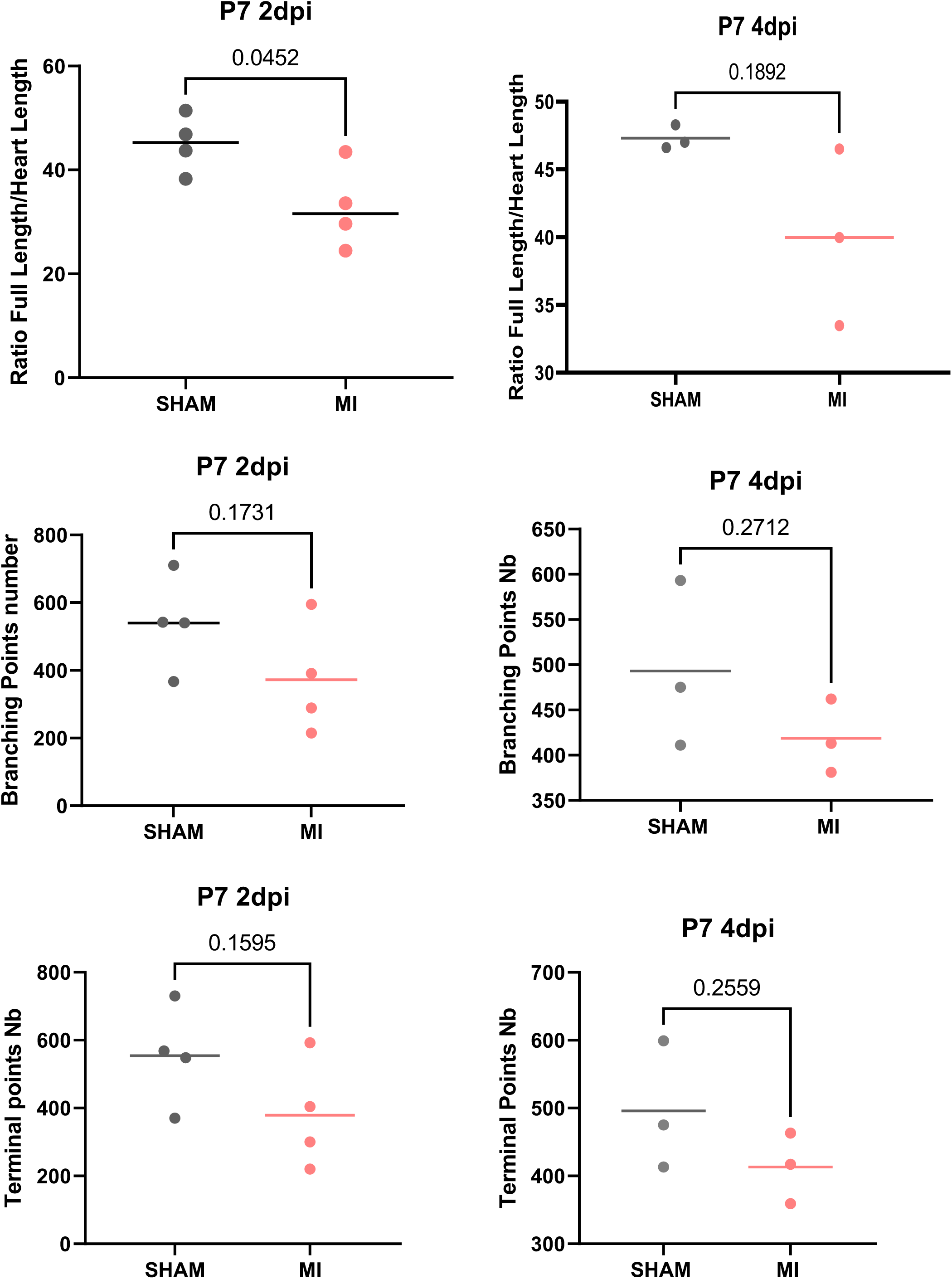
Comparison of the arterial tree density (total filament length) and complexity (branch points and end points) between sham- and MIP7-operated hearts at 2 and 4 days post-injury. (t test, p value above each graph, N: each dot corresponds to one heart)

**Supplementary figure 4:**
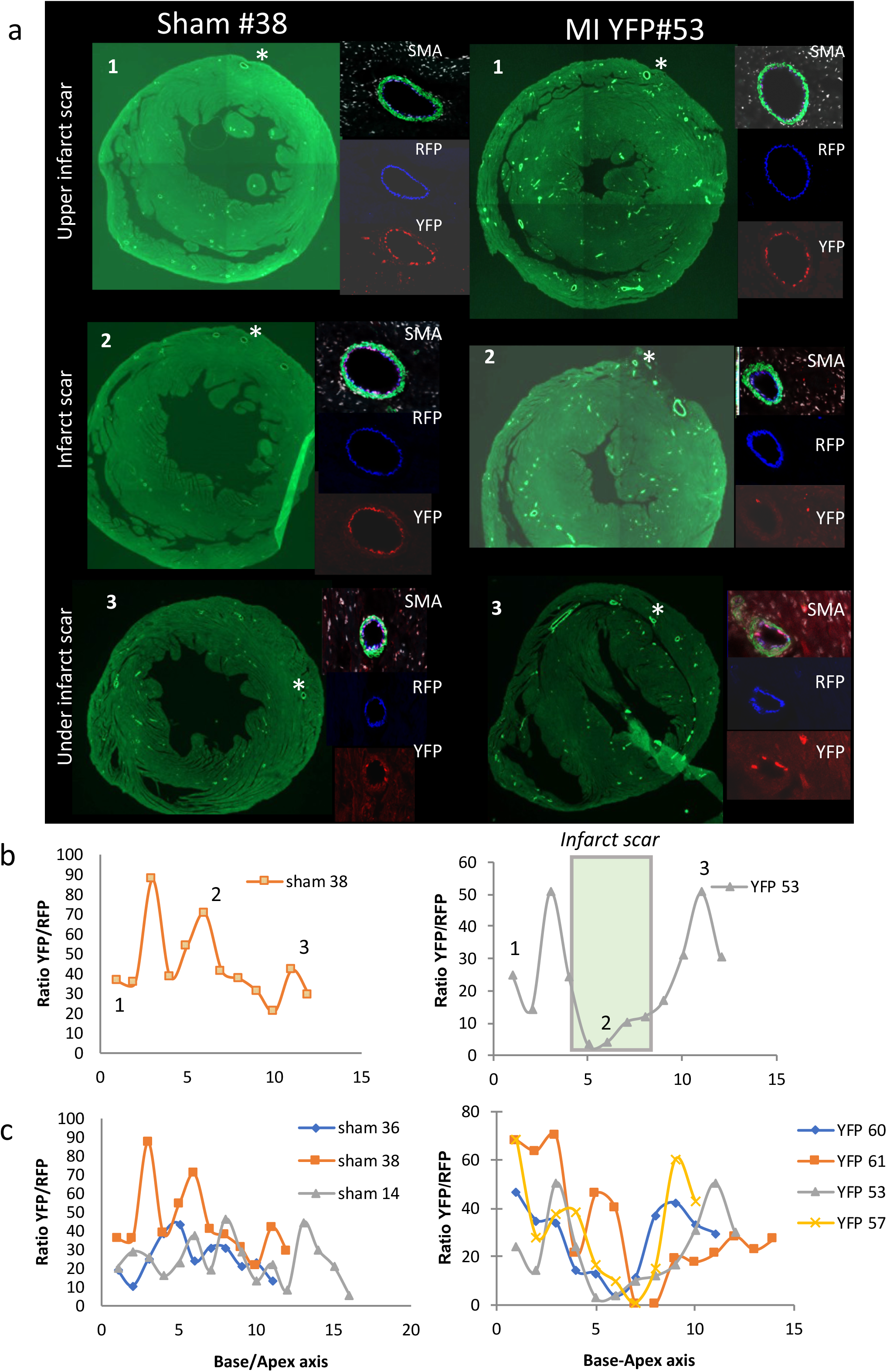
Genetic tracing of the embryonic Cx40-lineage (E14.5) during cardiac regeneration. **a-** Transverse sections of left ventricle from sham- and MIP1-operated *Cx40-CreERT2/RYFP* hearts stained for SMA (smooth muscle cells), RFP (arterial endothelial cells) and GFP (Cx40-lineage) at three levels below the ligation points. **b-** Graphs representative of the YFP/RFP proportion from sham- and MIP1-operated *Cx40-CreERT2/RYFP* hearts in successive sections from the ligation point (0) to the apex (15). **c-** Graphs representative of the YFP/RFP proportion from 3 sham- and 3 MIP1-operated *Cx40-CreERT2/RYFP* hearts in successive sections from the ligation point (0) to the apex (15).

**Supplementary figure 5:**
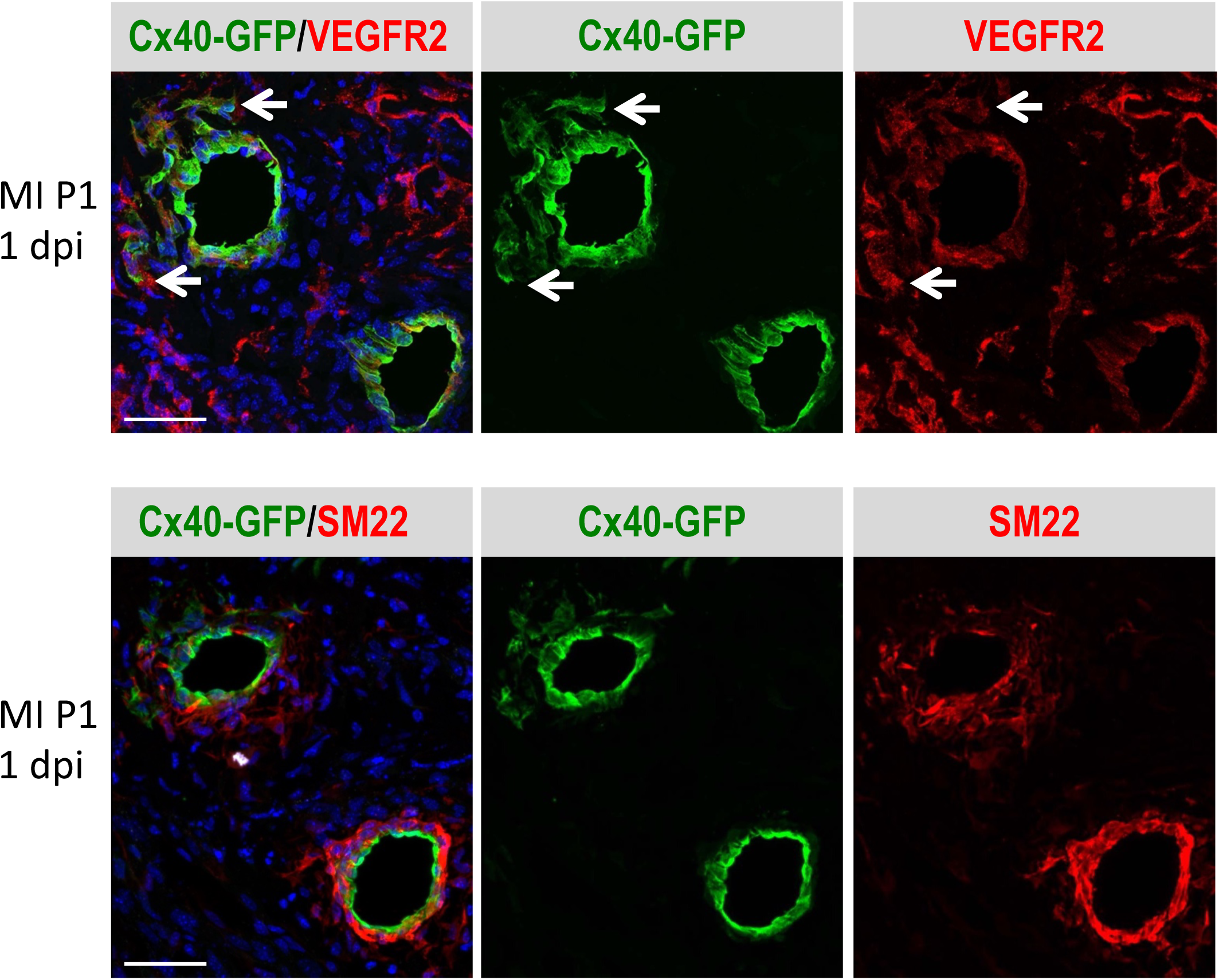
High magnification of the Left coronary artery below the ligation point on transverse section stained by Cx40-GFP (arterial cells) and VEGFR2 (capillaries and endothelial progenitors) or SM22 (smooth muscle cells). Arrows indicate the presence of isolated arterial cells leaving the Left coronary artery one day after injury from a MIP1 heart. Scale bar= 50µm.

## References

[1] A. Cannata, H. Ali, G. Sinagra, M. Giacca, Gene Therapy for the Heart Lessons Learned and Future Perspectives, Circ Res 126(10) (2020) 1394–1414.

[2] I.E. Lupu, S. De Val, N. Smart, Coronary vessel formation in development and disease: mechanisms and insights for therapy, Nat Rev Cardiol 17(12) (2020) 790–806.

[3] T. Su, G. Stanley, R. Sinha, G. D’Amato, S. Das, S. Rhee, A.H. Chang, A. Poduri, B. Raftrey, T.T. Dinh, W.A. Roper, G. Li, K.E. Quinn, K.M. Caron, S. Wu, L. Miquerol, E.C. Butcher, I. Weissman, S. Quake, K. Red-Horse, Single-cell analysis of early progenitor cells that build coronary arteries, Nature 559(7714) (2018) 356–362.

[4] B. Sharma, A. Chang, K. Red-Horse, Coronary Artery Development: Progenitor Cells and Differentiation Pathways, Annu Rev Physiol 79 (2017) 1–19.

[5] L. He, Q. Liu, T. Hu, X. Huang, H. Zhang, X. Tian, Y. Yan, L. Wang, Y. Huang, L. Miquerol, J.D. Wythe, B. Zhou, Genetic lineage tracing discloses arteriogenesis as the main mechanism for collateral growth in the mouse heart, Cardiovasc Res 109(3) (2016) 419–30.

[6] L. Miquerol, J. Thireau, P. Bideaux, R. Sturny, S. Richard, R.G. Kelly, Endothelial plasticity drives arterial remodeling within the endocardium after myocardial infarction, Circ Res 116(11) (2015) 1765–71.

[7] K.N. Dube, T.M. Thomas, S. Munshaw, M. Rohling, P.R. Riley, N. Smart, Recapitulation of developmental mechanisms to revascularize the ischemic heart, JCI Insight 2(22) (2017).

[8] J. Tang, H. Zhang, L. He, X. Huang, Y. Li, W. Pu, W. Yu, L. Zhang, D. Cai, K.O. Lui, B. Zhou, Genetic Fate Mapping Defines the Vascular Potential of Endocardial Cells in the Adult Heart, Circ Res 122(7) (2018) 984–993.

[9] C. Seiler, The human coronary collateral circulation, Eur J Clin Invest 40(5) (2010) 465–76.

[10] P. Meier, S. Gloekler, R. Zbinden, S. Beckh, S.F. de Marchi, S. Zbinden, K. Wustmann, M. Billinger, R. Vogel, S. Cook, P. Wenaweser, M. Togni, S. Windecker, B. Meier, C. Seiler, Beneficial effect of recruitable collaterals: a 10-year follow-up study in patients with stable coronary artery disease undergoing quantitative collateral measurements, Circulation 116(9) (2007) 975–83.

[11] H. Zhang, J.E. Faber, De-novo collateral formation following acute myocardial infarction: Dependence on CCR2(+) bone marrow cells, J Mol Cell Cardiol 87 (2015) 4–16.

[12] D. Chalothorn, J.A. Clayton, H. Zhang, D. Pomp, J.E. Faber, Collateral density, remodeling, and VEGF-A expression differ widely between mouse strains, Physiol Genomics 30(2) (2007) 179–91.

[13] S. Anbazhakan, P.E. Rios Coronado, A.N.L. Sy-Quia, L.W. Seow, A.M. Hands, M. Zhao, M.L. Dong, M.R. Pfaller, Z.A. Amir, B.C. Raftrey, C.K. Cook, G. D’Amato, X. Fan, I.M. Williams, S.K. Jha, D. Bernstein, K. Nieman, A.M. Pasca, A.L. Marsden, K.R. Horse, Blood flow modeling reveals improved collateral artery performance during the regenerative period in mammalian hearts, Nat Cardiovasc Res 1(8) (2022) 775–790.

[14] S. Das, A.B. Goldstone, H. Wang, J. Farry, G. D’Amato, M.J. Paulsen, A. Eskandari, C.E. Hironaka, R. Phansalkar, B. Sharma, S. Rhee, E.A. Shamskhou, D. Agalliu, V. de Jesus Perez, Y.J. Woo, K. Red-Horse, A Unique Collateral Artery Development Program Promotes Neonatal Heart Regeneration, Cell 176(5) (2019) 1128–1142 e18.

[15] E. Tzahor, K.D. Poss, Cardiac regeneration strategies: Staying young at heart, Science 356(6342) (2017) 1035–1039.

[16] B.J. Haubner, J. Schneider, U. Schweigmann, T. Schuetz, W. Dichtl, C. Velik-Salchner, J.I. Stein, J.M. Penninger, Functional Recovery of a Human Neonatal Heart After Severe Myocardial Infarction, Circ Res 118(2) (2016) 216–21.

[17] R. Marin-Juez, M. Marass, S. Gauvrit, A. Rossi, S.L. Lai, S.C. Materna, B.L. Black, D.Y. Stainier, Fast revascularization of the injured area is essential to support zebrafish heart regeneration, Proc Natl Acad Sci U S A 113(40) (2016) 11237–11242.

[18] T. Kocijan, M. Rehman, A. Colliva, E. Groppa, M. Leban, S. Vodret, N. Volf, G. Zucca, A. Cappelletto, G.M. Piperno, L. Zentilin, M. Giacca, F. Benvenuti, B. Zhou, R.H. Adams, S. Zacchigna, Genetic lineage tracing reveals poor angiogenic potential of cardiac endothelial cells, Cardiovasc Res 117(1) (2021) 256–270.

[19] H. Kolesova, M. Capek, B. Radochova, J. Janacek, D. Sedmera, Comparison of different tissue clearing methods and 3D imaging techniques for visualization of GFP-expressing mouse embryos and embryonic hearts, Histochem Cell Biol 146(2) (2016) 141–52.

[20] L. Miquerol, S. Meysen, M. Mangoni, P. Bois, H.V. van Rijen, P. Abran, H. Jongsma, J. Nargeot, D. Gros, Architectural and functional asymmetry of the His-Purkinje system of the murine heart, Cardiovasc Res 63(1) (2004) 77–86.

[21] S. Beyer, R.G. Kelly, L. Miquerol, Inducible Cx40-Cre expression in the cardiac conduction system and arterial endothelial cells, Genesis 49(2) (2011) 83–91.

[22] S. Srinivas, T. Watanabe, C.S. Lin, C.M. William, Y. Tanabe, T.M. Jessell, F. Costantini, Cre reporter strains produced by targeted insertion of EYFP and ECFP into the ROSA26 locus, BMC Dev Biol 1 (2001) 4.

[23] L. Madisen, T.A. Zwingman, S.M. Sunkin, S.W. Oh, H.A. Zariwala, H. Gu, L.L. Ng, R.D. Palmiter, M.J. Hawrylycz, A.R. Jones, E.S. Lein, H. Zeng, A robust and high-throughput Cre reporting and characterization system for the whole mouse brain, Nat Neurosci 13(1) (2010) 133–40.

[24] E.A. Susaki, K. Tainaka, D. Perrin, H. Yukinaga, A. Kuno, H.R. Ueda, Advanced CUBIC protocols for whole-brain and whole-body clearing and imaging, Nat Protoc 10(11) (2015) 1709–27.

[25] G. D’Amato, R. Phansalkar, J.A. Naftaly, X. Fan, Z.A. Amir, P.E. Rios Coronado, D.O. Cowley, K.E. Quinn, B. Sharma, K.M. Caron, A. Vigilante, K. Red-Horse, Endocardium-to-coronary artery differentiation during heart development and regeneration involves sequential roles of Bmp2 and Cxcl12/Cxcr4, Dev Cell 57(22) (2022) 2517–2532 e6.

[26] G. Heusch, Myocardial ischaemia-reperfusion injury and cardioprotection in perspective, Nature Reviews Cardiology 17 (2020) 773–789.

[27] C. Ryan, E.W. Gertz, Fistula from coronary arteries to left ventricle after myocardial infarction, Br Heart J 39(10) (1977) 1147–9.

[28] S.A. Said, R.H. Schiphorst, R. Derksen, L.J. Wagenaar, Coronary-cameral fistulas in adults: Acquired types (second of two parts), World J Cardiol 5(12) (2013) 484–94.

[29] C. Seiler, M. Stoller, B. Pitt, P. Meier, The human coronary collateral circulation: development and clinical importance, Eur Heart J 34(34) (2013) 2674–82.

[30] M. Potente, H. Gerhardt, P. Carmeliet, Basic and therapeutic aspects of angiogenesis, Cell 146(6) (2011) 873–87.

[31] J. Chen, D.K. Ceholski, L. Liang, K. Fish, R.J. Hajjar, Variability in coronary artery anatomy affects consistency of cardiac damage after myocardial infarction in mice, Am J Physiol Heart Circ Physiol 313(2) (2017) H275–H282.

[32] A.H. Chang, B.C. Raftrey, G. D’Amato, V.N. Surya, A. Poduri, H.I. Chen, A.B. Goldstone, J. Woo, G.G. Fuller, A.R. Dunn, K. Red-Horse, DACH1 stimulates shear stress-guided endothelial cell migration and coronary artery growth through the CXCL12-CXCR4 signaling axis, Genes Dev 31(13) (2017) 1308–1324.

[33] I. Buschmann, A. Pries, B. Styp-Rekowska, P. Hillmeister, L. Loufrani, D. Henrion, Y. Shi, A. Duelsner, I. Hoefer, N. Gatzke, H. Wang, K. Lehmann, L. Ulm, Z. Ritter, P. Hauff, R. Hlushchuk, V. Djonov, T. van Veen, F. le Noble, Pulsatile shear and Gja5 modulate arterial identity and remodeling events during flow-driven arteriogenesis, Development 137(13) (2010) 2187–96.

[34] G. Arolkar, S.K. Kumar, H. Wang, K.M. Gonzalez, S. Kumar, B. Bishnoi, P.E. Rios Coronado, Y.J. Woo, K. Red-Horse, S. Das, Dedifferentiation and Proliferation of Artery Endothelial Cells Drive Coronary Collateral Development in Mice, Arterioscler Thromb Vasc Biol 43(8) (2023) 1455–1477.

